# The RAP2.12 and RAP2.3 factors act downstream of LRR-MAL Receptor Kinases in *Arabidopsis* pollen-stigma interactions

**DOI:** 10.64898/2026.08.24.746730

**Authors:** Stephen J. Bordeleau, Younyoung Lee, Marcus A. Samuel, Daphne R. Goring

**Affiliations:** Department of Cell & Systems Biology, University of Toronto, Toronto ON, Canada M5S 3B2; Department of Biological Sciences, University of Calgary, Calgary AB, Canada T2N 1N4

**Keywords:** ERFVIIs, Pollen-pistil interactions, Pollen tube callose plugs, Receptor kinases, RKF1, RAP2.12, RAP2.3

## Abstract

*Arabidopsis Leucine-Rich Repeat-Malectin Receptor Kinase* (*LRR-MAL RK*) genes have been previously implicated in the early stages of pollen-pistil interactions to support compatible pollen. One member, *Receptor Kinase in Flowers 1* (*RKF1*), has been associated with roles in the stigma to support pollen hydration as well as pollen tube growth. To better understand RKF1’s function in these processes, a yeast two-hybrid screen was conducted with the RKF1 cytosolic kinase domain. Two positive interactors identified from this screen were the Group VII Ethylene Response Factors (ERFVIIs), RELATED TO APETALA 2.12 (RAP2.12) and RAP2.3. Their putative roles in pollen-pistil interactions were investigated using the quintuple *erfvii* mutant, and novel pistil-mediated pollen tube callose deposition phenotypes were uncovered during the pollen tube growth stage. Loss of seven *LRR-MAL RKs* including *RKF1* in the pistil was previously found to cause an unusual phenotype where shorter callose plugs were deposited in wildtype pollen tubes compared to that seen in wildtype Col-0 pistils. Contrary to this, wildtype pollen tubes growing through the quintuple *erfvii* mutant pistil deposited callose plugs that were more elongated than that seen in wildtype Col-0 pistils. Further analyses with the *proteolysis 6* (*prt6*) mutant and *RAP2.12* rescue constructs were consistent with these phenotypes providing support that *RKF1* is a negative regulator of *RAP2.12* and *RAP2.3* in the pistil during pollen tube growth.

## Introduction

In *Arabidopsis*, the process of fertilization begins with self-pollination where pollen grains are deposited on the stigmatic papillae located at the top the pistil. This contact leads to a rapid release of water from the stigmatic papilla to the pollen for hydration. This is then followed by the emergence of a pollen tube that enters into the stigmatic papilla cell wall and grows down to its base. The pollen tube then enters the reproductive tract, growing between the cells of the solid tissue in the stigma and style until it reaches the transmitting tract in the ovule where cells surrounded by extracellular matrix material are loosely spaced out. Here the pollen tube is guided to an unfertilized ovule where sperm cells are delivered to the female gametophyte for fertilization. Receptors kinases play a prominent role throughout this process, starting from pollen contact with a stigmatic papilla to the end where the pollen tube delivers sperm cells to an unfertilized ovule (reviewed in (Zhong et al., 2025;Zhang et al., 2026).

Previously, we have uncovered roles for *Arabidopsis* Leucine-Rich Repeat-Malectin Receptor Kinases (LRR-MAL RKs, also known as LRR-VIII-2 RKs) in the stigma and pistil reproductive tract to promote pollen hydration and support pollen tube growth (Lee and Goring, 2021;Lee et al., 2024). Specifically, *Receptor Kinase in Flowers 1* (*RKF1*), along with its three tandemly-linked paralogues, *RKFL1-3*, are required in the stigma to promote efficient pollen hydration. Three additional *LRR-MAL RK* family members, L*ysM RLK1-interacting Kinase 1* (*LIK1*; (Le et al., 2014), *Remorin-Interacting Receptor 1* (*RIR1*; (Abel et al., 2021) and *Nematode-Induced LRR-RLK 2* (*NILR2*; (Mendy et al., 2017) were also investigated, but the additional loss of these three *LRR-MAL RKs* failed to compound this pollen hydration phenotype, indicating the loss of the *RKF1* gene cluster was the primary cause for reduced pollen hydration (Lee and Goring, 2021;Lee et al., 2024). Interestingly, the knockout of all seven *LRR-MAL RKs* (*RKF1, RKFL1-3, LIK1, RIR1* and *NILR2*) uncovered a role in the pistil in promoting pollen tube growth through the stigma and style tissues. Wildtype pollen tubes growing through this septuple *LRR-MAL RK* mutant pistil displayed shorter callose plugs in the pollen tubes (Lee and Goring, 2021;Lee et al., 2024). Callose plugs are deposited in the pollen tube to partition the cytoplasm, along with the vegetative cell and two sperm cells, to the front of the growing pollen tube (Adhikari et al., 2020;Kapoor and Geitmann, 2023). This callose plug phenotype in the wildtype pollen tubes was thought to result from morphological changes to the pollen tube as it grew invasively through the mutant stigma and style tissues lacking the seven *LRR-MAL RKs* (Lee et al., 2024). Additionally, the *RKF1* gene cluster combined with *SOMATIC EMBRYOGENESIS RECEPTOR KINASE 1* (*SERK1*) and *BRI1-ASSOCIATED RECEPTOR KINASE 1* (*BAK1/SERK3*) was found to be required in the pistil to support pollen tube growth as wildtype pollen tubes displayed shorter lengths when growing through the *rkfΔ serk1 bak1* sextuple mutant pistils (Lee and Goring, 2021).

To get further insight into how the *LRR-MAL RKs* are functioning in the upper pistil to influence *Arabidopsis* pollen-pistil dynamics, a yeast two-hybrid screen was conducted with the RKF1 cytosolic kinase domain to identify potential downstream signalling proteins. From this screen, two members of the Group VII Ethylene Response Factors (ERFVIIs), RELATED TO APETALA 2.12 (RAP2.12) and RAP2.3, were identified. *Arabidopsis* RAP2.12 and RAP2.3 have well-characterized roles in regulating hypoxia responsive genes in response to hypoxia, and more recently *Arabidopsis* RAP2.12 was implicated in regulating developmental events in hypoxic niches (reviewed in (Renziehausen et al., 2024;Gibbs et al., 2025;Wang et al., 2025). Interestingly in the absence of hypoxia, RAP2.12 and RAP2.3 can be sequestered at the plasma membrane for protection against proteasomal degradation by the N-degron pathway, and their activity is also regulated by phosphorylation from several different protein kinases (reviewed in (Gibbs et al., 2025;Wang et al., 2025). These different features suggest that RAP2.12 and RAP2.3 could be valid interactors of RKF1 and were investigated further for potential functions in *Arabidopsis* pollen-pistil interactions.

## Materials and methods

### Plant materials and growth conditions

All seeds were stratified in water at 4°C in the dark for a minimum of 5 days prior to sowing directly onto Sunshine #1 soil. Soil was supplemented with Plant Prod All-Purpose 20-20-20 fertilizer for growth. All plants were grown in growth chambers set to a 16h/8hr light/dark cycle at 23°C with the relative humidity kept under 50%. Three sets of *Arabidopsis* Col-0 mutants were used in this study: the quintuple *rap2.12-1 rap2.2-1 rap2.3-1 hre1-1 hre2-1* mutant (*erfvii* mutant (Abbas et al., 2015;Leon et al., 2020), the *prt6-1* (SAIL 1278_H11) and *prt6-5* (SALK_051088C) mutants (Graciet et al., 2009), and the septuple *rkfΔ-1 nilr2rir1Δ-1 lik1-5* mutant (Lee et al., 2024). The mutant genotypes were confirmed for all plants used in this study by PCR genotyping with primers listed in Supplementary Table S4.

### Yeast two-hybrid analyses

Hybrigenics Services (France) conducted two screens using the *ATFO* yeast two-hybrid library consisting of random-primed cDNA synthesized from *Arabidopsis* Col-0 RNA, extracted from flower buds and open flowers (Supplementary Tables S1 and S2). The RKF1 cytosolic kinase domain (starting immediately after the transmembrane domain) was used as the bait, and each screen tested 123 million interactions. The first screen was conducted on plates with 10 mM 3-aminotriazol (to reduce background), and 285 positive clones were picked and sequenced (Supplementary Table S1). For the second screen, 3-aminotriazol was omitted, and all yeast colonies from the screen were pooled (∼ 5000-6000). Plasmid DNA was then extracted from the pooled yeast colonies and sequenced by Illumina sequencing (MiSeq V3-300 x 2 paired end sequencing, University of Toronto CAGEF facility), and the data was analyzed by Hybrigenics Services to produce a more extensive list of potential interactors (Supplementary Table S2). Among the analytic data provided for each screen was the predicted interaction region for each candidate (See Supplementary Tables S1 and S2 for more details).

Candidate interactors were then selected to validate interactions by yeast two-hybrid screening (Fig. 1; Supplementary Table S3). Col-0 RNA was extracted from whole leaves using the RNeasy® Plant Mini Kit (Qiagen) and used to synthesize cDNA with SuperScript™ IV Reverse Transcriptase (Invitrogen). The cDNA was used to PCR amplify regions corresponding to either full length proteins or predicted interacting domains for the candidate interactors (Supplementary Table S3; primers listed in Supplementary Table S4). The PCR products were tagged with attB1 and attB2 sites to gateway clone into the pDONR207 entry plasmid using BP-Clonase II (Life Technologies). Entry clones were then incubated with LR-Clonase II (Life Technologies) to clone the cDNAs into the yeast two-hybrid destination vector, pJG4-5. Plasmid DNA for these final clones were transformed into the EGY48 yeast strain following a small-scale library transformation protocol (DupLEX-A Yeast Two-Hybrid System Application Guide, OriGene). Pairwise interactions were tested for each candidate interactors against the RKF1 and LIK1 cytosolic kinase domains using yeast two-hybrid mating assays, and positive interactions were identified by the activation of the *LacZ* reporter leading to the formation of blue yeast colonies as described in (Bastedo et al., 2019;Lee and Goring, 2021).

**Fig. 1.**
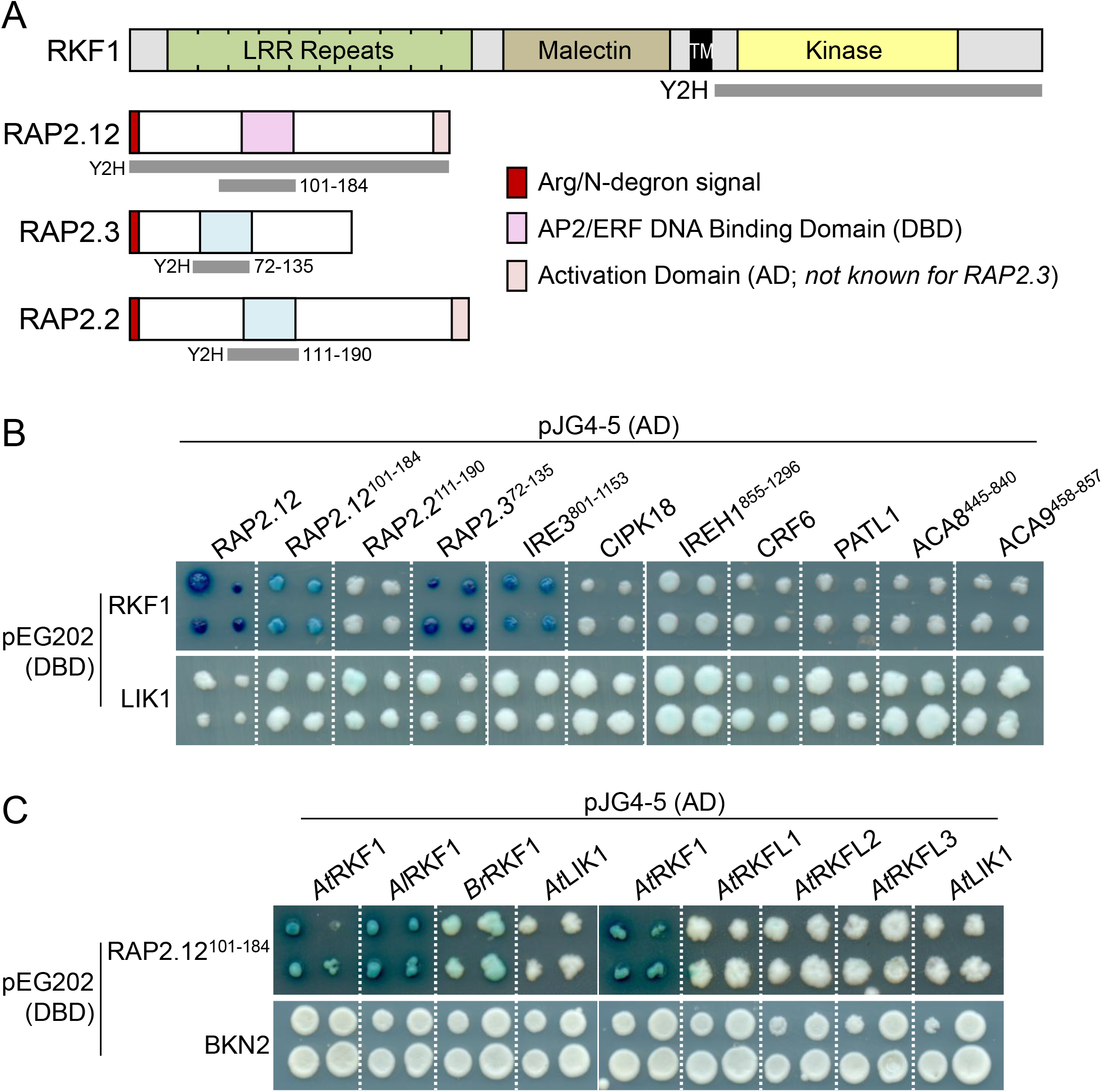
Pairwise Y2H interaction screen uncovers new RKF1 interactors. (A) Diagrams of the predicted protein domains in RKF1 and three ERFVII factors. The regions tested in the Y2H screen are shown in gray. AD: activation domain. DBD: DNA Binding Domain. (B) The RKF1 cytosolic kinase domain was screened against candidate interactors using full length proteins or predicted interaction domains derived from the Y2H library screens (Tables S1-S3). The domains for the ERFVII factors are shown in A. IRE3^801-1153^ and IREH1^855-1296^ are kinase domains; ACA8^445-840^ and ACA9^458-857^ are cytosolic loop 3 domains (Table S3). The LIK1 cytosolic kinase domain was used as a negative control to rule out auto-activation by candidate interactors. (C) RAP2.12^101-184^ was screened against different RKF1-related cytosolic kinase domains and interacted with RKF1 orthologues from *Arabidopsis lyrata* and *Brassica rapa*, but not with the RKF1 paralogs, AtRKFL1-3. The BKN2 kinase was used as a negative control to rule out auto-activation by the different cytosolic kinase domains. For all receptor kinases, the cytosolic kinase domains start immediately after the predicted transmembrane domains (Lee and Goring, 2021).

### Recombinant protein production, in vitro kinase assay, and phosphoprotein gel staining

The coding region corresponding to the RKF1 cytosolic kinase domain (starting immediately after the transmembrane domain) was synthesized and cloned as a GST fusion in pET-42a(+), while the full length coding sequences for RAP2.12 and RAP2.3 were synthesized and cloned as His-tagged fusions in pET-28a(+) (GenScript). These constructs were transformed into *E. coli* BL21 (DE3) cells to produce the fusion proteins, and expression was induced by adding 0.4 mM isopropyl β-D-1-thiogalactopyranoside at 16°C for 16 hours. Harvested bacterial cells were resuspended in lysis buffer, lysed by sonication, and the lysates were then incubated with the corresponding resin overnight at 4°C with gentle agitation. For the His-tagged RAP2.12 and RAP2.3, the lysis buffer contained 20 mM Tris-HCl (pH 8.0), 150 mM NaCl, 20 mM imidazole, 5% glycerol, 0.5% Tween 20, 1 mM PMSF, and 1 cOmplete™ tablet (Protease Inhibitor Cocktail, Roche), and the lysates were incubated with Ni-NTA resin (Qiagen). For GST-RKF1, the lysis buffer contained 50 mM HEPES (pH 7.5), 150 mM NaCl, 3 mM DTT, 0.05% Triton X-100, and 1 cOmplete™ tablet, and the lysate was incubated with Glutathione Sepharose 4B resin (GE Healthcare). The fusion proteins were purified from these resins using sequential washing steps to remove non-specific protein interactions, following by elution of the recombinant proteins from the resins. The fusion proteins were verified by western blot analysis, using anti-His antibodies (RAP2.12 and RAP2.3) or anti-GST antibodies (RKF1). The GST:RKF1 and 6xHis:RAP2.3 proteins were similar in size to their predictions (GST:RKF1, 70 kDa and 6xHis:RAP2.3, 30 kDa) while 6xHis:RAP2.12 migrated higher than its predicted size of 42 kDa, but this was previously observed by (Gasch et al., 2016) as well.

In vitro kinase assays were performed using purified GST-RKF1, His-RAP2.12, and His-RAP2.3 proteins with a 2× kinase reaction buffer (0.25 M HEPES pH 7.5, 5 mM MnCl₂, 0.5 mM dithiothreitol and 50 μM ATP) and incubated at 30°C for 90 minutes. Reactions were terminated by adding 10 μL of 4× SDS loading buffer, followed by boiling at 98°C for 10 minutes. Kinase reaction products were separated on a 10% SDS-PAGE gel, and phosphoprotein detection was performed using Pro-Q Diamond Phosphoprotein Stain following the manufacturer’s protocol (Thermo Fisher Scientific). Phosphoproteins in the gel were imaged using an iBright Imager (Invitrogen), and the gel was stained with Coomassie Brilliant Blue (CBB) to visualize the proteins.

### RAP2.12 rescue constructs and Agrobacterium mediated plant transformation

To generate plant lines expressing *RAP2.12* in the stigmas of Col-0 and *erfvii* plants, two forms of RAP2.12 were synthesized with the stigma-specific *Arabidopsis SLR1* promoter (*AtS1*; (Dwyer et al., 1994) and cloned into the pORE3 transformation vector as HindIII-MluI fragments (GeneArt, Thermo Fisher Scientific). The constructs either had the wildtype RAP2.12 gene sequence or the Δ13RAP2.12 gene sequence which has the first 13-amino acids constituting its N-degron tag removed (Licausi et al., 2011). Sequences are shown in Supplementary Fig. S1. A third construct with the *Arabidopsis SLR1* promoter fused to *GUS* was also synthesized, cloned into pORE3 as a HindIII-SpeI fragment and used to verify stigma-specific expression (Supplementary Fig. S2). Wildtype (Col-0) and erfvii plants were transformed with these transformation vectors through Agrobacterium-mediated floral dipping, and plants were dipped twice with a 7-day interval to maximize the transformation efficiency (Clough and Bent, 1998). Seeds from dipped plants were collected and stratified for a week prior to sowing onto soil. Germinated seedlings were sprayed 3 times at 2-day intervals with a BASTA herbicide solution (6:1000 dilution of a 10 g/L solution) to select for positive transformants. Transformants were confirmed through PCR genotyping of genomic DNA for the BASTA gene.

### Pollen hydration assays and aniline blue staining of pollinated pistils

Pollination assays were conducted as described in (Lee et al., 2020) and all pollinations used wildtype Col-0 pollen grains to pollinate wildtype Col-0, mutant or transgenic pistils. For both the pollen hydration assays and aniline blue staining of pollinated pistils, stage 12 flower buds were emasculated and in the morning of the following day, the now stage 13 pistils were hand-pollinated by lightly brushing a single anther over the stigmatic papillae of the receptive pistil. For pollen hydration assays, images were captured of the pollinated stigmas at 0 minutes and 10 minutes, immediately following pollen deposition (post-pollination). The diameters of the same pollen grains were measured from the captured images at 0 minutes and 10 minutes as a proxy for pollen hydration. Images were captured using a Nikon SMZ800 stereo zoom microscope with a DS-Fi1 color camera, and measurements were taken for 30 pollen grains for each line using the NIS Element software (3 stigmas each with 10 pollen grains/stigma randomly sampled).

For aniline blue staining of pollen tubes in pollinated pistils, hand-pollinated pistils were left for 2 hours to allow for pollen tube growth through the stigma and style. After 2 hours, the pollinated pistils were collected and fixed in a 3:1 ethanol - glacial acetic acid solution for at least 30 minutes at room temperature. Fixed pistils were washed three times with water and then incubated in a 0.5M NaOH solution for approximately 40 minutes at 60°C. Pistils were again washed three times with water and placed in a 0.1% decolorized Aniline Blue stain and left for 30 minutes at room temperature or overnight in the dark at 4°C. For imaging, the stained pistils were mounted in water, some pressure was added to the coverslip to slightly flatten the stigma and imaged at 10X magnification on a Zeiss Axioskop2Plus fluorescent microscope. Both brightfield and UV (aniline blue) images were captured using a Lumenera Infinity3S color camera. To measure callose plug lengths, callose plugs were randomly sampled from 3-5 aniline blue-stained pistils, and lengths were measured using the INFINITY ANALYZE software.

## Results

### Verifying candidate RKF1 Interactors from the yeast two-hybrid screens

To identify candidate intracellular binding partners for RKF1, two yeast two-hybrid screens were conducted using a random-primed cDNA library made from RNA extracted from flower buds and open flowers (Hybrigenics Services, see methods for more details). From the first screen, 285 positive clones were sequenced and RAP2.12, an ERFVII transcription factor, was the top interactor in the set with 171 of the 285 clones corresponding to RAP2.12 (Supplementary Table S1). In the second screen, all the positive colonies were pooled for sequencing which producing a more extensive list of potential interactors (Supplementary Table S2). Several criteria were used to develop a shorter list of potential candidate interactors for verification from the second screen. This included signaling-related protein annotations, gene expression profiles in female reproductive tissues, predicted protein subcellular localization in the cytosol or at the plasma membrane, and the predicted interaction regions to be cytosolic domains. From this, ten candidates were selected for re-testing in the yeast two-hybrid system (Fig. 1, Supplementary Table S3).

RAP2.12 was again a top interactor in the second screen, and its AP2/ERF DNA binding domain was predicted to be the minimal interaction domain in both screens. Another ERFVII, RAP2.3, was also uncovered in the second screen with its AP2/ERF DNA binding domain included in the predicted minimal interaction domain. The remaining three ERFVII members, RAP2.2, HYPOXIA RESPONSE ELEMENT 1 (HRE1) and HRE2 (Gibbs et al., 2011), did not come up in this screen, but RAP2.2 was also tested based on its strong expression profile in pistil tissues (Supplementary Table S3; (Klepikova et al., 2016). Of the five ERFVII members, RAP2.12 and RAP2.3 are particularly notable as potential RKF1-interactors because they have been found to be sequestered at the plasma membrane under normal conditions and regulated by several different protein kinases (reviewed in (Gibbs et al., 2025;Wang et al., 2025). Several other members of the AP2/ERF superfamily (Ma et al., 2024) also came up in the second screen, and one member, Cytokinin Response Factor 6 (CRF6) was included for verification as it was found in both screens and was predicted to interact with the RKF1 kinase domain via its AP2/ERF DNA binding domain as well (Supplementary Tables S1-S3).

Of this set of candidates, only RAP2.12 and RAP2.3 were verified as interactors of RKF1 in the yeast two hybrid analysis (Fig. 1). Both the full length RAP2.12 clone and the RAP2.12^101-184^ clone consisting of the AP2/ERF domain were positive interactors (Fig 1. A-B). Similarly, the RAP2.3^72-135^ AP2/ERF domain clone also was a positive interactor. These clones interacted with RFK1 but not with the negative control which consisted of the cytosolic kinase domain of another LRR-MAL RK, LIK1 (Fig. 1B). The other two candidates containing AP2/ERF domains, RAP2.2 and CRF6, did not interact with RKF1 suggesting there may be some specificity to the RAP2.12 and RAP2.3 AP2/ERF domains interacting with RKF1 (Fig. 1B). Interestingly, the interaction between RKF1 and RAP2.12^101-184^ was found to be exclusive to RKF1 among the tandemly linked RKF1 gene cluster, with no interactions detected for RKFL1, 2 and 3 (Fig. 1C). The RKF1-RAP2.12 interaction was also found to be conserved with orthologues from other Brassicaceae species as positive interactions were seen with cytosolic kinase domains from *A. lyrata* RKF1 and *Brassica rapa* RKF1 (Fig. 1C).

Using the criteria described above, the remaining set of candidate interactors selected for re-testing in the yeast two-hybrid assay included two proteins kinases (Incomplete Root Hair Elongation 3 [IRE3] and CBL-Interacting Protein Kinase 18 [CIPK18]), two calcium pumps (Autoinhibited Ca^2+^-ATPase 8 [ACA8] and ACA9), and a membrane trafficking protein (PATELLIN-1 [PATL1]) (Supplementary Table S3). With ACA8 and ACA9 being multi-pass transmembrane proteins, the predicted interaction domain, cytoplasmic loop 3 (Supplementary Table S3), was tested for both candidates. Of this set, only IRE3 displayed a positive interaction with RKF1 (Fig. 1B). IRE3 is an AGC kinase in the *AGC-Other* subfamily, and along with its homologue IREH1, is proposed to regulate microtubule organization in the growing root as knockout mutants displayed disordered microtubules and skewed roots (Bogre et al., 2003;Yue et al., 2019). Despite IREH1 being closely related to IRE3, it did not interact with RKF1 (Fig 1B). In the following sections, we focused on the RKF1-RAP2.12/RAP2.3 interactions, and the RKF1-IRE3 interaction will be followed up in another study.

Since RKF1’s kinase domain was shown to be a functional serine-threonine kinase (Takahashi et al., 1998), we investigated whether RKF1 could phosphorylate RAP2.12 and RAP2.3. Fusion proteins consisting of a GST:RKF1 cytosolic kinase domain fusion and His-tagged fusions for RAP2.12 and RAP2.3 were produced and added to *in vitro* kinase assays. The kinase reactions were then separated on a SDS-PAGE gel, followed by staining with the Pro-Q diamond phosphoprotein gel stain to detect phosphoproteins (Fig 2). RKF1 was confirmed to catalyze autophosphorylation (Fig. 2, lane 1) and found to phosphorylate both RAP2.12 and RAP2.3 (Fig. 2, lanes 4-5). Overall, the yeast-2-hybrid and phosphorylation assays demonstrating a clear relationship between RKF1 and RAP2.12/RAP2.3, and so this prompted us to further investigate potential biological functions of these ERFVIIs during pollen-pistil interactions.

**Fig. 2.**
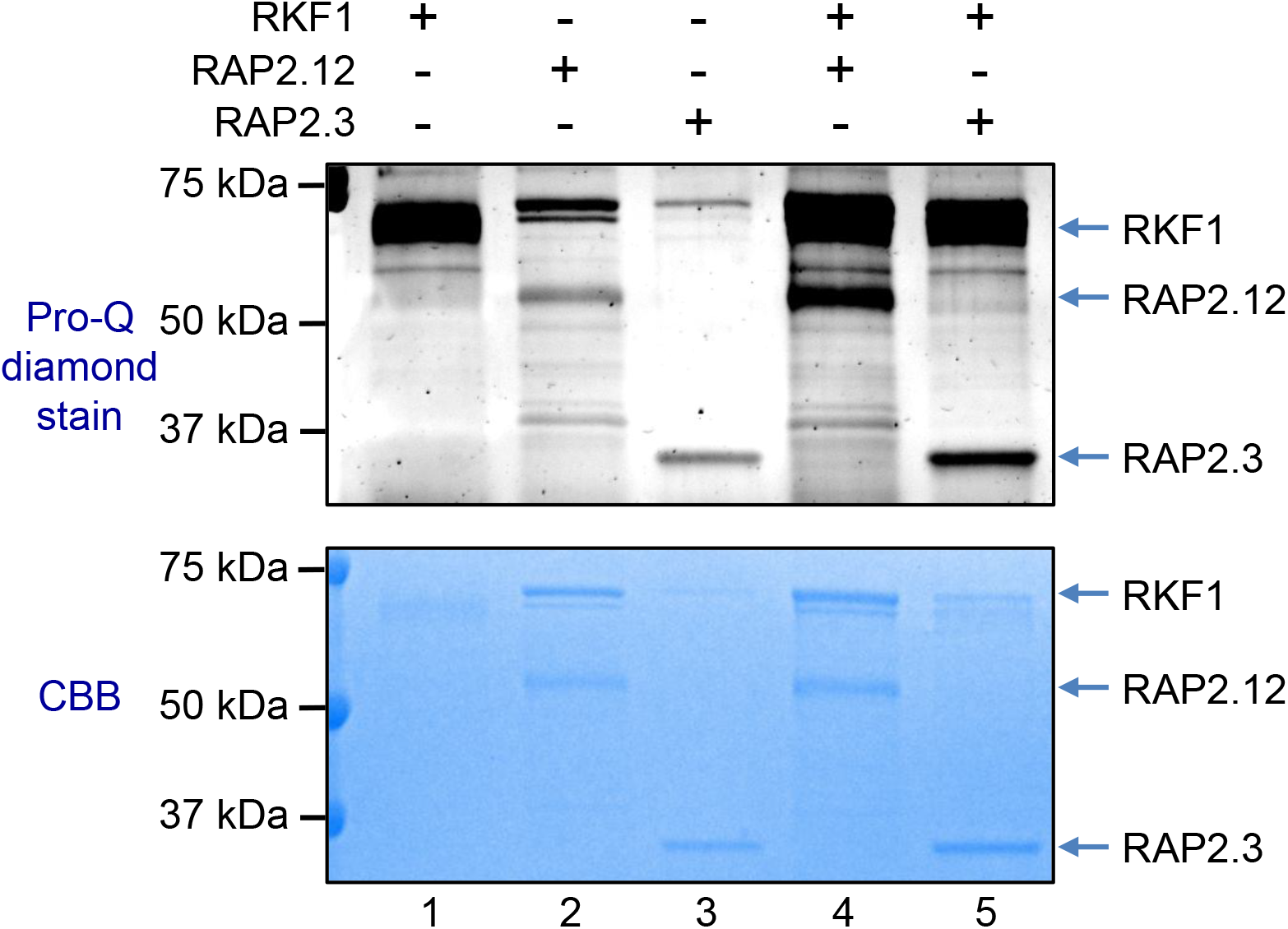
RKF1 phosphorylation of RAP2.12 and RAP2.3. *In vitro* kinase assays were performed using purified fusion proteins to test for RKF1 autophosphorylation and phosphorylation of RAP2.12 and RAP2.3 by RKF1. For RKF1, GST was fused to the N-terminus of the RKF1 cytosolic kinase domain, start immediately after the predicted transmembrane domain. For RAP2.12 and RAP2.3, the 6xHis tag was fused to the N-termini of these full-length proteins. These fusion proteins were added to *in vitro* kinase assays, and the reactions were run on a 12% SDS-gel, stained with Pro-Q Diamond Phosphoprotein gel stain, followed by CBB staining to visualize protein bands. Predicted protein sizes are as follows: GST:RKF1, 70 kDa; 6xHis:RAP2.12, 42 kDa*; 6xHis:RAP2.3, 30 kDa. *As seen here, RAP2.12 was previously found to migrate higher than its predicted size (Gasch et al., 2016).

### Assessing pollen-pistil interactions in the quintuple ERFVII mutant

The *RKF1* gene cluster has been previously implicated in the stigma to support pollen hydration and in the upper pistil to support pollen tube growth (with the latter role involving other LRR MAL RKs and SERK1 and BAK1; (Lee and Goring, 2021;Lee et al., 2024). To assess if the ERFVIIs have a role in the pistil to support wildtype pollen during the earlier pollen-pistil interaction stages, a well-characterized quintuple *rap2.12-1 rap2.2-1 rap2.3-1 hre1-1 hre2-1* mutant was used (hereafter *erfvii* mutant; (Marin-de la Rosa et al., 2014;Abbas et al., 2015;Gibbs et al., 2018;Shukla et al., 2019;Leon et al., 2020;Zubrycka et al., 2023;Eysholdt-Derzso et al., 2024). It is important to note that while five ERFVII family members were knocked out in the *erfvii* mutant, only RAP2.12, RAP2.2 and RAP2.3 are expressed at higher levels in the stigma (Supplementary Table S3). First, the ability of *erfvii* mutant stigmas to support wildtype Col-0 pollen hydration was assessed by hand-pollinating wildtype Col-0 and *erfvii* mutant stigmas and measuring wildtype Col-0 pollen grain diameters at 0- and 10-minutes post-pollination. At 10-minutes post-pollination, there were no significant differences in wildtype Col-0 pollen hydration on wildtype Col-0 stigmas versus *erfvii* stigmas (Fig. 3A). This indicates that that *ERFVIIs* are not involved in the stigma to support pollen hydration.

**Fig. 3.**
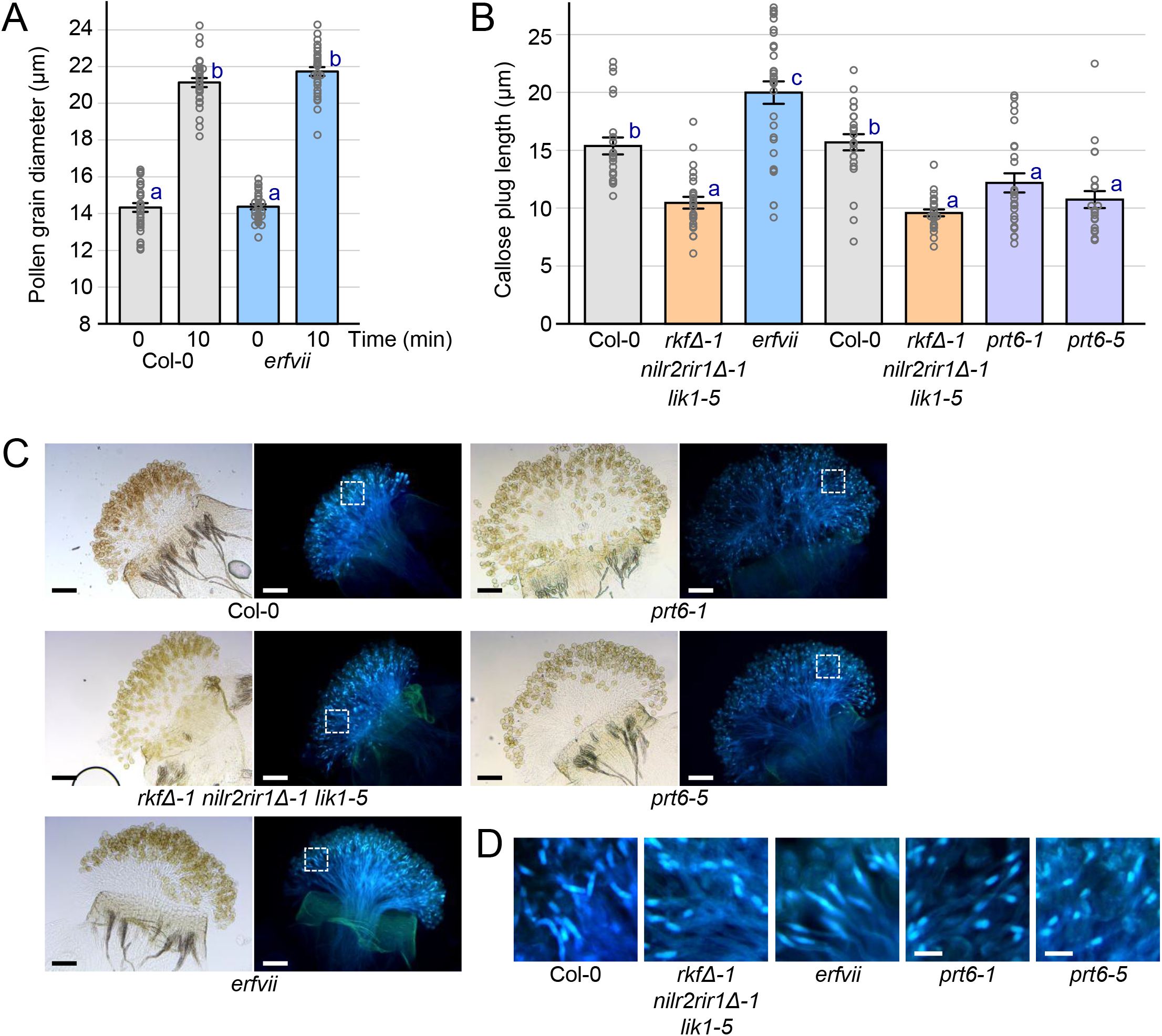
Wildtype Col-0 pollen tubes display altered callose plug lengths at 2 hours post-pollination on stigmas from the *erfvii* and *prt6* mutants. (A) Quantification of wildtype Col-0 pollen hydration at 0- and 10-minutes post-pollination on stigmas from Col-0 and the *erfvii* mutant. Pollen grain diameters were measured as a proxy for pollen hydration (Lee *et al*., 2020). n = 30 pollen grains per line. (B) Callose plug lengths in wildtype Col-0 pollen tubes growing in stigmas from wildtype Col-0 and mutants at 2 hours post-pollination. n = 23-31 callose plugs per line. (C) Representative images of wildtype Col-0 pollen tubes with callose plugs in stigmas from Col-0 and mutants at 2 hours post-pollination. Brightfield images (left) and aniline blue stained images (right) are shown for each sample. Scale bar = 100µm. (D) Close up views of callose plugs in wildtype Col-0 pollen tubes growing in Col-0 and mutant stigmas at 2 hours post-pollination. White boxes in (C) outline the regions shown here. Scale bar = 20µm. Data in (A, B) are shown as bar graphs +/- SE with all data points displayed. Letters represent statistically different groupings of P<0.05 based on a one-way ANOVA with a Tukey HSD post-hoc test. The *erfvii* mutant = the quintuple *rap2.12-1 rap2.2-1 rap2.3-1 hre1-1 hre2-1* mutant (Gibbs *et al*., 2014). The *rkfΔ-1 nilr2rir1Δ-1 lik1-5 mutant* serves as a control for shorter callose plugs as reported in Lee *et al*. (2024).

Next, we examined wildtype Col-0 pollen tube growth through the *erfvii* mutant stigma/style tissues. Prior work had uncovered an interesting phenotype presenting in wildtype pollen tubes growing through the stigma and style of a septuple *LRR-MAL RK* mutant pistil. The loss of seven *LRR-MAL RKs* (*RKF1 RKFL1-3 NILR2 RIR1 LIK1*) in the mutant pistil (*rkfΔ-1 nilr2rir1Δ-1 lik1-5*) resulted in significantly shorter callose plugs deposited in wildtype pollen tubes compared to those deposited in wildtype pistils (Lee et al., 2024). This phenotype presented at 2-hours post-pollination when the pollen tubes were growing through the stigma/style tissues (Fig. 3B-D). To determine if this stage of pollen tube growth was affected in the *erfvii* mutant, the *erfvii* pistils were hand-pollinated with wildtype pollen grains and harvested at 2-hours post-pollination for aniline blue staining of pollen tubes and imaging. Remarkably, the callose plugs deposited in wildtype pollen tubes growing through *erfvii* pistils were found to display the opposite phenotype of the septuple *rkfΔ-1 nilr2rir1Δ-1 lik1-5* mutant pistils. The wildtype pollen tubes in the *erfvii* stigmas contained elongated callose plugs in comparison to wildtype pollen tubes presents in the wildtype Col-0 stigmas and the *rkfΔ-1 nilr2rir1Δ-1 lik1-5* mutant stigmas (Fig. 3B-D). Measuring the lengths of these callose plugs revealed a statistically significant difference between those deposited in the wildtype pollen tubes in the stigmas from the *erfvii* mutant compared to wildtype Col-0 and the *rkfΔ-1 nilr2rir1Δ-1 lik1-5* mutant (Fig. 3B). The wildtype pollen tubes also displayed the previously observed shorter callose plugs when growing through the *rkfΔ-1 nilr2rir1Δ-1 lik1-5* mutant stigmas compared to wildtype Col-0 stigmas (Fig. 3B). Despite these changes in the callose plug lengths, the overall reproductive success was not affected as normal seed set was observed in the *erfvii* and *rkfΔ-1 nilr2rir1Δ-1 lik1-5* mutants.

Based on the opposite callose plug length phenotypes in wildtype pollen tubes growing through the *erfvii* mutant stigmas versus the *rkfΔ-1 nilr2rir1Δ-1 lik1-5* mutant stigmas, we hypothesized that the LRR-MAL RKs may be negatively regulators of the ERFVIIs in the stigma/style tissues. To test this idea, we investigated the potential impact of having increased ERFVIIs in stigmas on wildtype pollen tubes. In the absence of hypoxia, the five ERFVIIs are regulated by the N-degron pathway where in the final step, the Proteolysis 6 (PRT6) E3-ubiquitin ligase target the ERFVIIs for proteasomal degradation. (Gibbs et al., 2011;Licausi et al., 2011;Gibbs et al., 2025). It has been shown that the ERFVIIs are more stable in the *prt6* mutant and so we tested two independent *prt6* T-DNA mutant pistils in the 2-hour wildtype pollen tube growth assays. Interestingly, callose plugs in the wildtype pollen tubes growing through *prt6* mutant stigmas displayed the same shorter callose plug phenotype as seen for wildtype pollen tubes in the *rkfΔ-1 nilr2rir1Δ-1 lik1-5* mutant stigmas (Fig. 3C-D) with no significant difference (Fig 3B). Given this observed phenotype is opposite to that observed in wildtype pollen tubes in the *erfvii* mutant stigmas, this supports a role for LRR-MAL RKs, much like PRT6, for negatively regulating the ERFVIIs in some manner in the stigma/style tissues.

### The stigma-specific erfvii mutant rescue of elongated callose plugs in wildtype Col-0 pollen tubes

To confirm the elongated callose plug phenotype in wildtype Col-0 pollen tubes was associated with the loss of *ERFVIIs* in the *erfvii* mutant stigmas, the stigma-specific *Arabidopsis SLR1* promoter (Dwyer et al., 1994) was used to express *RAP2.12* in wildtype Col-0 and *erfvii* mutant plants. Two different *RAP2.12* constructs were tested in these transgenic plants (Supplementary Fig. S1): (i) *SLR1p:RAP2.12* which includes the N-terminal degron sequence that would target RAP2.12 for degradation by PRT6 in the absence of hypoxia and (ii) *SLR1p:Δ13RAP2.12* where the RAP2.12 N-terminal degron (13 amino acids) was omitted to allow for RAP2.12 accumulation in the absence of hypoxia (Gibbs et al., 2011;Licausi et al., 2011;Gibbs et al., 2025). Three independent transgenic lines from each transformation were analyzed in detail for rescue of the elongated callose plugs in wildtype pollen tubes (Fig. 4-5, Supplementary Fig. S3). A third construct, placing the *GUS* reporter downstream of the *SLR1* promoter, was transformed into Col-0 plants to confirm that the *SLR1* promoter directed expression strictly to the stigma in the flower (Supplementary Fig. S1-S2).

**Fig. 4.**
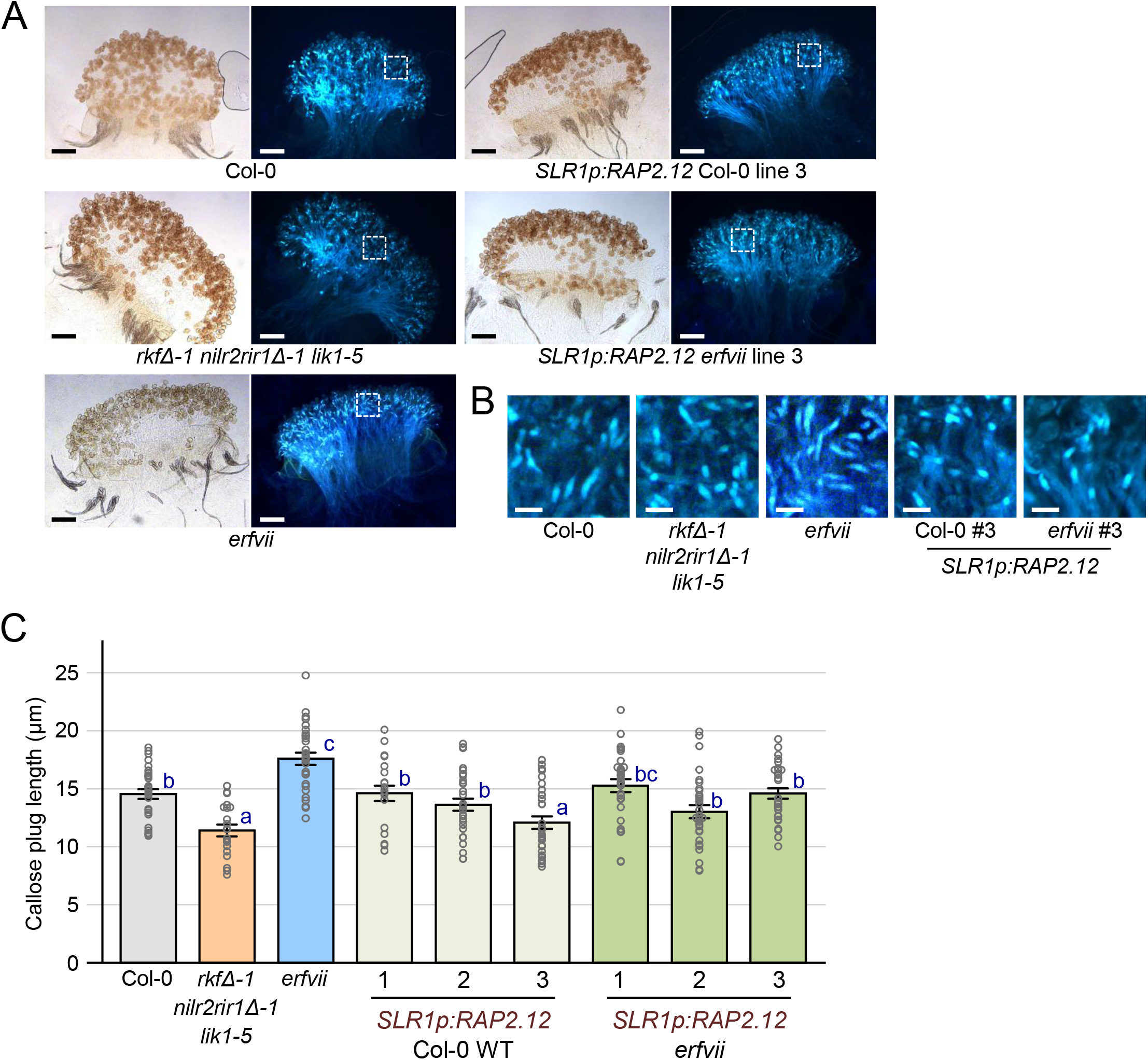
The expression of *RAP2.12* in *erfvii* mutant stigmas rescues the longer callose plug phenotype in wildtype Col-0 pollen tubes at 2 hours post-pollination. (A) Representative images of wildtype Col-0 pollen tubes with callose plugs in stigmas from Col-0 and the *SLR1p:RAP2.12* rescue lines at 2 hours post-pollination. Brightfield images (left) and aniline blue stained images (right) are shown for each sample. Scale bar = 100µm. See Supplementary Fig. S2 for additional images. (B) Close up views of callose plugs in wildtype Col-0 pollen tubes growing in stigmas from Col-0 and *SLR1p:RAP2.12* rescue lines at 2 hours post-pollination. White boxes in (A) outline the regions shown here. Scale bar = 20µm. (C) Callose plug lengths in wildtype Col-0 pollen tubes growing in stigmas from Col-0 and *SLR1p:RAP2.12* rescue lines at 2 hours post-pollination. n = 20-30 callose plugs per line. Data is shown as a bar graph +/- SE with all data points displayed. Letters represent statistically different groupings of P<0.05 based on a one-way ANOVA with a Tukey HSD post-hoc test. The *erfvii* mutant = the quintuple *rap2.12-1 rap2.2-1 rap2.3-1 hre1-1 hre2-1* mutant (Gibbs *et al*., 2014). The *rkfΔ-1 nilr2rir1Δ-1 lik1-5 mutant* serves as a control for shorter callose plugs as reported in Lee *et al*. (2024).

**Fig. 5.**
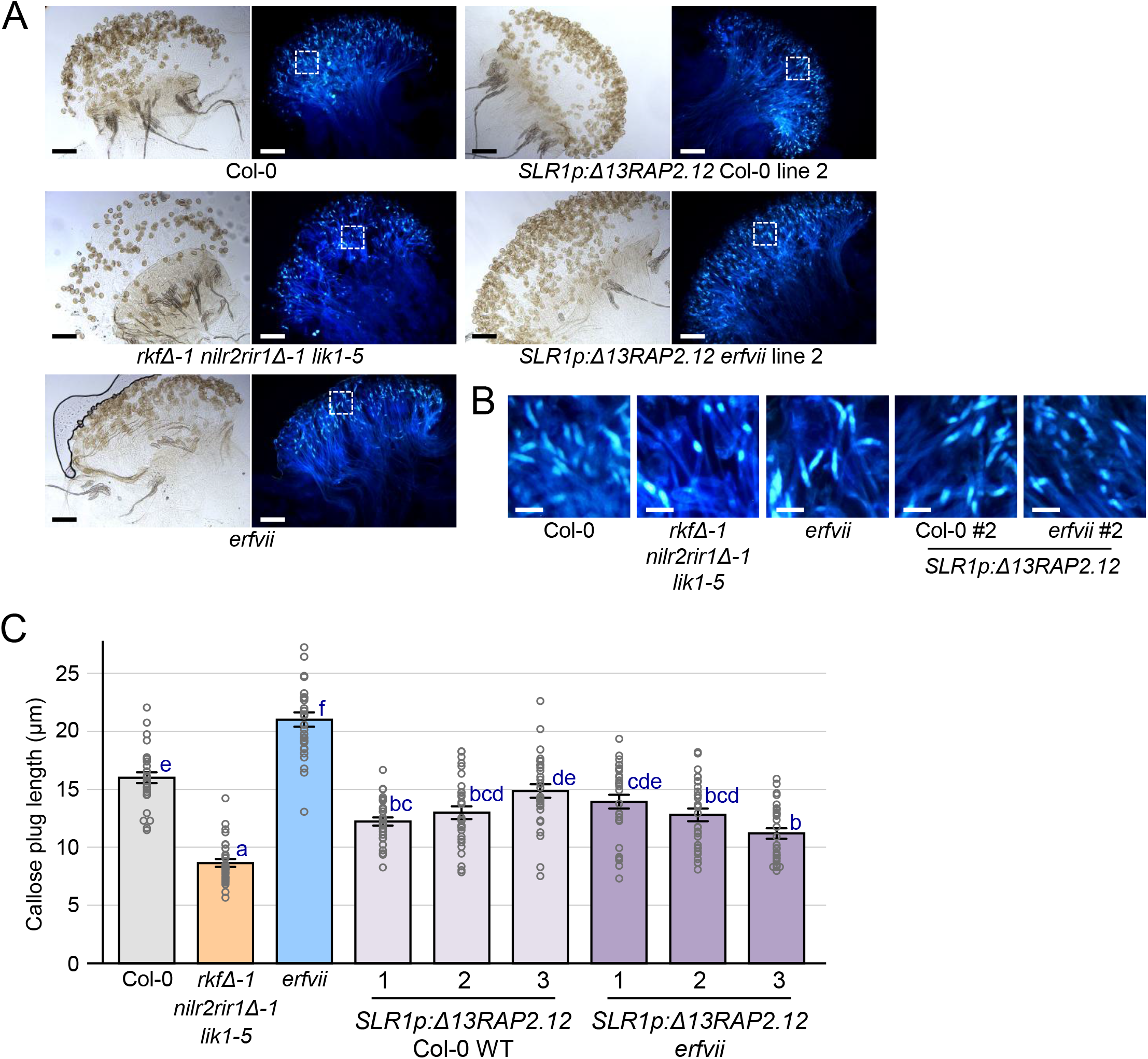
The expression of *Δ13RAP2.12* in *erfvii* mutant stigmas rescues the longer callose plug phenotype in wildtype Col-0 pollen tubes at 2 hours post-pollination. (A) Representative images of wildtype Col-0 pollen tubes with callose plugs in stigmas from Col-0 and the *SLR1p:Δ13RAP2.12* rescue lines at 2 hours post-pollination. Brightfield images (left) and aniline blue stained images (right) are shown for each sample. Scale bar = 100µm. See Supplementary Fig. S2 for additional images. (B) Close up views of callose plugs in wildtype Col-0 pollen tubes growing in stigmas from Col-0 and the *SLR1p:Δ13RAP2.12* rescue lines at 2 hours post-pollination. White boxes in (A) outline the regions shown here. Scale bar = 20µm. (C) Callose plug lengths in wildtype Col-0 pollen tubes growing in stigmas from Col-0 and the *SLR1p:Δ13RAP2.12* rescue lines at 2 hours post-pollination. n = 30 callose plugs per line. Data is shown as a bar graph +/- SE with all data points displayed. Letters represent statistically different groupings of P<0.05 based on a one-way ANOVA with a Tukey HSD post-hoc test. The *erfvii* mutant = the quintuple *rap2.12-1 rap2.2-1 rap2.3-1 hre1-1 hre2-1* mutant (Gibbs *et al*., 2014). The *rkfΔ-1 nilr2rir1Δ-1 lik1-5 mutant* serves as a control for shorter callose plugs as reported in Lee *et al*. (2024).

To evaluate the *RAP2.12* transgenic plants, pistils were hand-pollinated with wildtype Col-0 pollen and left for 2-hours prior to aniline blue staining and imaging. Overall, the expression of *RAP2.12* and *Δ13RAP2.12* in *erfvii* stigmas was able to rescue the elongated callose plug phenotype in wildtype pollen tubes. This was seen by wildtype pollen tubes containing shorter callose plugs at 2-hours post-pollination compared to those growing through the *erfvii* stigmas (Fig. 4-5, Supplementary Fig. S3). The expression of *RAP2.12* in *erfvii* stigmas restored the callose plugs in the wildtype pollen tubes to lengths that were the equivalent to that observed in wildtype Col-0 stigmas (Fig. 4C). Interestingly though, the expression of *Δ13RAP2.12* in *erfvii* stigmas reduced the callose plug lengths in the wildtype pollen tubes to significantly shorter lengths in two transgenic lines (#2 and #3) than that observed in wildtype Col-0 stigmas (Fig. 5C). This may suggest that having more RAP2.12 activity in the stigma is associated with wildtype pollen tubes depositing shorter callose plugs. This trend was also observed for wildtype pollen tubes growing through wildtype Col-0 stigmas (Fig. 4-5, Supplementary Fig. S3). For stigmas from one of three *RAP2.12* transgenic Col-0 lines and two of the three *Δ13RAP2.12* transgenic Col-0 lines, the callose plug lengths in the wildtype pollen tubes had significantly shorter lengths compared to that observed in wildtype Col-0 stigmas (Fig. 4C and 5C). Despite these decreased callose plug lengths in the wildtype pollen tubes in the rescue lines, the callose plugs were generally longer that that observed in wildtype pollen tubes in the *rkfΔ-1 nilr2rir1Δ-1 lik1-5* mutant stigmas (Fig. 4C and 5C). Overall, the expression of *RAP2.12/Δ13RAP2.12* in the *erfvii* mutant stigmas successfully rescued the elongated callose plug phenotype in wildtype Col-0 pollen tubes growing through *erfvii* mutant stigmas.

## Discussion

Previously, the *Arabidopsis RKF1 Leucine-Rich Repeat-Malectin Receptor Kinase* (*LRR-MAL RK*) was shown to play a role in the pistil to support the early stages of pollen-pistil interactions (Lee and Goring, 2021;Lee et al., 2024). To further understand how RKF1 functions in this role, potential RFK1 intracellular binding partners were identified from a yeast two-hybrid screen (Fig. 1-2). Two of these RKF1 interactors, RAP2.12 and RAP2.3, were further analyzed, and we uncovered new roles for these Group VII Ethylene Response Factors (ERFVIIs) in the pistil to support wildtype pollen. There were two phenotypes associated with the loss of RKF1 that were examined in this study. First, stigmas from a CRISPR deletion mutant of the *Arabidopsis RKF1* gene cluster (quadruple *rkfΔ* mutant) supported reduced hydration of wildtype pollen (Lee and Goring, 2021). However, this phenotype was not observed for wildtype pollen placed on *Arabidopsis erfvii* mutant stigmas indicating that *RAP2.12* and *RAP2.3* are not involved at this stage (Fig. 3). Second, the loss of the *RKF1* gene cluster, along with three other *Arabidopsis LRR-MAL RKs* (septuple *rkfΔ-1 nilr2rir1Δ-1 lik1-5* mutant) resulted in wildtype pollen tubes depositing shorter callose plugs when growing through the stigmas of these mutant pistils (Lee et al., 2024). Interestingly here, we observed that the callose plugs in wildtype pollen tubes growing through the *erfvii* mutant stigmas were longer than that observed in wildtype pollen tubes growing through wildtype Col-0 stigmas (Fig. 3). These opposite phenotypes suggest opposing genetic functions for these *Arabidopsis LRR-MAL RKs* and *ERFVIIs* in female reproductive tissues during the early stages of pollen tube growth.

The *Arabidopsis* ERFVIIs are key players in promoting survival during environmental hypoxic stresses, and *Arabidopsis* RAP2.12 has also been implicated in regulating developmental outcomes in endogenous hypoxic niches. Of the five *Arabidopsis* ERFVIIs, RAP2.12 and RAP2.3 are particularly interesting as their activity can be regulated in distinct ways. In the absence of hypoxia, the *Arabidopsis* ERFVIIs are regulated by the N-degron pathway and subject to proteasomal degradation, but RAP2.12 and RAP2.3 can also be sequestered at the plasma membrane through interactions with Acyl-CoA Binding Proteins (ACBPs) for protection against proteasomal degradation (Schmidt and van Dongen, 2019). RAP2.12 is also regulated by several protein kinases during environmental hypoxic stresses. Typically, these protein kinases are positive regulators that phosphorylate RAP2.12 to increase protein stability and transcriptional activity during hypoxia. These include the Hydraulic Conductivity of Root 1 (HCR1) Raf-like kinase, the Target of Rapamycin (TOR) kinase, the MPK3 and MPK6 MAP kinases, and the CPK12 calcium-dependent protein kinase (Shahzad et al., 2016;Zhou et al., 2022;Kunkowska et al., 2023;Zhao et al., 2025). However, a recent study uncovered MPK9 as a negative regulator of RAP2.12 where MPK9 phosphorylation of RAP2.12 was connected to reduced RAP2.12 stability during hypoxia (Zhou et al., 2026). Here, we proposed that RKF1’s negative regulatory relationship with RAP2.12 and RAP2.3 could also be connected to their stability in the stigma as a result of phosphorylation. Using an *in vitro* kinase assay, we found that RKF1 will phosphorylate RAP2.12 and RAP2.3 (Fig. 2), but whether this might affect RAP2.12/RAP2.3 stability is not known.

To test if RAP2.12 and RAP2.3 stability in the stigma is connected to the pollen tube callose plug phenotype, we used the *prt6* mutant which knocks out the N-degron pathway resulting in increased accumulation of the ERFVIIs (Gibbs et al., 2011;Licausi et al., 2011;Gibbs et al., 2025). Stigmas from the *prt6* mutant would be predicted to promote a similar callose plug phenotype in wildtype pollen tubes as that observed for the septuple *LRR-MAL RK* (*rkfΔ-1 nilr2rir1Δ-1 lik1-5*). Indeed, the loss of *PRT6* function in the stigma did result in the same shorter callose plug phenotype in the wildtype pollen (Fig. 3). Additionally, the expression of *RAP2.12* in the *erfvii* stigmas rescues the elongated callose plug phenotype in wildtype pollen tubes (Fig. 4-5). These findings suggest that RKF1 is potentially regulating RAP2.12 and RAP2.3 stability to maintain optimal levels. Zhou et al. (2026) proposed that the function of MPK9 phosphorylation of RAP2.12 was to “fine-tune” RAP2.12 levels during hypoxia, and RKF1 may have a similar function in the stigma transmitting tract. It will need to be investigated in future work if altered levels of these ERFVIIs (increased or decreased) are somehow changing the growth environment in the stigma which is turn alters the callose plug morphology in the wildtype pollen tubes.

The path of pollen tube starts with growth through the cell wall of the stigmatic papilla on the surface of the stigma and down to the base of the papilla. From there, the pollen tube grows between the closely packed cells of the stigma and style transmitting tissue down towards the ovary transmitting tissue that has a much looser organization of cells with abundant extracellular matrix material surrounding the cells (Ndinyanka Fabrice et al., 2017;Reimann et al., 2020;Robichaux and Wallace, 2021;Zhou et al., 2021;Kapoor and Geitmann, 2023). The changes observed in callose plug lengths in this study correspond to the stage where the pollen tubes are growing invasively through the densely packed cells of the stigma and style transmitting tissue and may reflect altered interactions between the wildtype pollen tubes and the mutant transmitting tissues. Seeing as *LRR-MAL RKs* are needed to support pollen tube growth through these tissues, perhaps the LRR-MAL RKs are perceiving some type of signal from the growing pollen tubes in this function. The LRR-MAL RK subfamily consists of 13 members that possess extracellular domains containing an LRR domain following by a malectin domain (reviewed in (Oelmuller et al., 2023;Vilchez-Pinto et al., 2026). One LRR-MAL RK member, IMPAIRED IN GLYCAN PERCEPTION1 (IGP1)/ CELLOOLIGOMER-RECEPTOR KINASE1 (CORK1), has been shown to bind to plant cell wall–derived damage-associated molecular patterns (DAMPs), specifically cellulose-derived oligosaccharides. IGP1/CORK1 has been proposed to be involved in cell wall integrity sensing and activates immune responses following binding of these cellooligomers (Tseng et al., 2022;Martin-Dacal et al., 2023;Fernández-Calvo et al., 2024). Structural studies have shown that IGP1/CORK1 bind to the cellooligomer, cellotriose, through the LRR domain but the cellotriose-binding pocket is specific to IGP1/CORK1 and not conserved in the other LRR-MAL RKs such as RKF1 (Jiménez-Sandoval et al., 2025;Sun et al., 2025). So whether RKF1 is sensing some cell wall derived oligosaccharides in the stigma arising from invasive pollen growth is one outstanding question stemming from this study as well as how the putative negative regulation of the ERFVIIs ties into this. Given RAP2.12’s involvement in developmental hypoxic niches (Shukla et al., 2019;Koo et al., 2024;Renziehausen et al., 2024), another area for future studies would be to investigate if a hypoxic niche is present in the dense stigma/style transmitting tissues and how this might impact pollen tube growth.

## Data availability

All data supporting the findings of this study are available within the paper and its Supplementary Data.

## Author contributions

SJB and DRG: conceptualization, formal analysis, and writing - original draft; SJB and YL: investigation; SJB, DRG, MAS, YL: visualization, and writing - review & editing; DRG and MAS: supervision and funding acquisition. All authors have read and approved the final manuscript.

## Acknowledgements

We thank work-study students (Minyoung Chung, Flower Tan, Daniel Haghi, Gary Chatha, Shreya Das) for their technical assistance. We are very grateful to Dr. Darrell Desveaux for providing the gateway-compatible yeast two-hybrid vectors, pJG4-5 and pEG202, and members of the Desveaux lab (Janis Cheng and Tamar Av-Shalom) for their assistance in the yeast two-hybrid screening. We are also very grateful to Dr. José León (Universidad Politécnica de Valencia) for the *erfvii* mutant seeds and the ABRC for the *prt6* mutant seeds (SAIL 1278_H11, SALK_051088C).

## Supplementary Data

**Figure S1.** Sequences for the SLR1 promoter-GUS and SLR1 promoter-RAP2.12 constructs. See methods for more information.

**Figure S2.** *Arabidopsis* SLR1 promoter:GUS transgenic lines showing stigma-specific GUS activity in the inflorescences.

**Figure S3.** The stigma expression of *RAP2.12* and *Δ13RAP2.12* rescues the longer callose plug phenotype in wildtype Col-0 pollen tubes at 2 hours post-pollination.

**Table S1 - List of RKF1 interacting proteins identified by yeast two-hybrid screen #1**

Sheet 1 – Legend

Sheet 2 - List of RKF1 interacting proteins identified by yeast two-hybrid screen #1

Sheet 3 - List of RKF1-RAP2.12 interactions from the yeast two-hybrid screen #1

**Table S2 - List of RKF1 interacting proteins identified by yeast two-hybrid screen #2**

Sheet 1 – Legend

Sheet 2 - Interactions

Sheet 3 - Background no sense

Sheet 4 - Background antisense

Sheet 5 - Top 150 IF clones annotated

**Table S3 - List of candidates re-tested in the yeast two-hybrid system**

Sheet 1 - List of candidates re-tested in the yeast two-hybrid system - interacting domain, type of protein and name, protein domains

Sheet 2 - TRAVA expression profiles of candidates re-tested in the yeast two-hybrid system

Sheet 3 - Predicted transmembrane, extracellular and cytoplasmic domains for ACA8 and ACA9

**Table S4.** Primers used in this study.

