## Supplementary material for "The RAP2.12 and RAP2.3 factors act downstream of LRR-MAL Receptor Kinases in *Arabidopsis* pollen-stigma interactions": Figures S1-S3

AtSLR1:GUS

AAGCTTTAACATCCGGTACGTTAATTGCAATTGTAAATCCAGTCTCACATGAAAACCGGATAGAATTGGATCTAACCCAATGAACCGGGAATCTAAATTAAAAATTAAAATTGCTCTAATA
AGACATGTTAATCCTAGAATGCTAATCATGCTTCCGTTATGTTTTCCCTCCGTGAACACTAAATGGAGAATCATCAACAATATGTTAAGGATGTGTTTGTTTGGGCTGCTTCGGCAACTTT
CGCTGCTTGTTAGATTGCGACTTATTGCGAGCATAGATGTCATGGAAGTAGAATTCGAAATACATCGAGAAGATTAAAATACATCGAACGTAACGATAAATGCATTTACGAGTGCCATAGC
ATGACTTTCTTGGGGACGAGGATCATGGATGAATAAATATACTTGTTCAACAACATATGTTATGTTGCACGAGCAAATTAATTTCACATGGTGACGTTATCGACTAATGACAGTAGTTTGT
TGAAGTGAATCAATGAGGTACTGAAAAGTCATAGAACGTGGAAAATTACATTTGTCTTGTCTGCTAGAAATAATTGTAATGAGTTTTGCAAATTTGAAAACCAACATGTGAAGAAATCTAT
AAATATAGGTTTTAAGAAACAAAGAAGAGCATAAGAAAGAAAGTCGTGAAG**ATG**TTACGTCCTGTAGAAACCCCAACCCGTGAAATCAAAAAACTCGACGGCCTGTGGGCATTCAGTCTGG
ATCGCGAAAACTGTGGAATTGATCAGCGTTGGTGGGAAAGCGCGTTACAAGAAAGCCGGGCAATTGCTGTGCCAGGCAGTTTTAACGATCAGTTCGCCGATGCAGATATTCGTAATTATGC
GGGCAACGTCTGGTATCAGCGCGAAGTCTTTATACCGAAAGGTTGGGCAGGCCAGCGTATCGTGCTGCGTTTCGATGCGGTCACTCATTACGGCAAAGTGTGGGTCAATAATCAGGAAGTG
ATGGAGCATCAGGGCGGCTATACGCCATTTGAAGCCGATGTCACGCCGTATGTTATTGCCGGGAAAAGTGTACGTATCACCGTTTGTGTGAACAACGAACTGAACTGGCAGACTATCCCGC
CGGGAATGGTGATTACCGACGAAAACGGCAAGAAAAAGCAGTCTTACTTCCATGATTTCTTTAACTATGCCGGAATCCATCGCAGCGTAATGCTCTACACCACGCCGAACACCTGGGTGGA
CGATATCACCGTGGTGACGCATGTCGCGCAAGACTGTAACCACGCGTCTGTTGACTGGCAGGTGGTGGCCAATGGTGATGTCAGCGTTGAACTGCGTGATGCGGATCAACAGGTGGTTGCA
ACTGGACAAGGCACTAGCGGGACTTTGCAAGTGGTGAATCCGCACCTCTGGCAACCGGGTGAAGGTTATCTCTATGAACTGTGCGTCACAGCCAAAAGCCAGACAGAGTGTGATATCTACC
CGCTTCGCGTCGGCATCCGGTCAGTGGCAGTGAAGGGCGAACAGTTCCTGATTAACCACAAACCGTTCTACTTTACTGGCTTTGGTCGTCATGAAGATGCGGACTTGCGTGGCAAAGGATT
CGATAACGTGCTGATGGTGCACGACCACGCATTAATGGACTGGATTGGGGCCAACTCCTACCGTACCTCGCATTACCCTTACGCTGAAGAGATGCTCGACTGGGCAGATGAACATGGCATC
GTGGTGATTGATGAAACTGCTGCTGTCGGCTTTAACCTCTCTTTAGGCATTGGTTTCGAAGCGGGCAACAAGCCGAAAGAACTGTACAGCGAAGAGGCAGTCAACGGGGAAACTCAGCAAG
CGCACTTACAGGCGATTAAAGAGCTGATAGCGCGTGACAAAAACCACCCAAGCGTGGTGATGTGGAGTATTGCCAACGAACCGGATACCCGTCCGCAAGGTGCACGGGAATATTTCGCGCC
ACTGGCGGAAGCAACGCGTAAACTCGACCCGACGCGTCCGATCACCTGCGTCAATGTAATGTTCTGCGACGCTCACACCGATACCATCAGCGATCTCTTTGATGTGCTGTGCCTGAACCGT
TATTACGGATGGTATGTCCAAAGCGGCGATTTGGAAACGGCAGAGAAGGTACTGGAAAAAGAACTTCTGGCCTGGCAGGAGAAACTGCATCAGCCGATTATCATCACCGAATACGGCGTGG
ATACGTTAGCCGGGCTGCACTCAATGTACACCGACATGTGGAGTGAAGAGTATCAGTGTGCATGGCTGGATATGTATCACCGCGTCTTTGATCGCGTCAGCGCCGTCGTCGGTGAACAGGT
ATGGAATTTCGCCGATTTTGCGACCTCGCAAGGCATATTGCGCGTTGGCGGTAACAAGAAAGGGATCTTCACTCGCGACCGCAAACCGAAGTCGGCGGCTTTTCTGCTGCAAAAACGCTGG
ACTGGCATGAACTTCGGTGAAAAACCGCAGCAGGGAGGCAAACAA**TGA**ACTAGT

AtSLR1:RAP2.12

AAGCTTTAACATCCGGTACGTTAATTGCAATTGTAAATCCAGTCTCACATGAAAACCGGATAGAATTGGATCTAACCCAATGAACCGGGAATCTAAATTAAAAATTAAAATTGCTCTAATA
AGACATGTTAATCCTAGAATGCTAATCATGCTTCCGTTATGTTTTCCCTCCGTGAACACTAAATGGAGAATCATCAACAATATGTTAAGGATGTGTTTGTTTGGGCTGCTTCGGCAACTTT
CGCTGCTTGTTAGATTGCGACTTATTGCGAGCATAGATGTCATGGAAGTAGAATTCGAAATACATCGAGAAGATTAAAATACATCGAACGTAACGATAAATGCATTTACGAGTGCCATAGC
ATGACTTTCTTGGGGACGAGGATCATGGATGAATAAATATACTTGTTCAACAACATATGTTATGTTGCACGAGCAAATTAATTTCACATGGTGACGTTATCGACTAATGACAGTAGTTTGT
TGAAGTGAATCAATGAGGTACTGAAAAGTCATAGAACGTGGAAAATTACATTTGTCTTGTCTGCTAGAAATAATTGTAATGAGTTTTGCAAATTTGAAAACCAACATGTGAAGAAATCTAT
AAATATAGGTTTTAAGAAACAAAGAAGAGCATAAGAAAGAAAGTCGTGAAG**ATG**TGTGGAGGAGCTATAATATCCGATTTCATTCCACCGCCGAGGTCTCGCCGTGTTACTAGCGAGTTTA
TTTGGCCGGATCTGAAGAAGAATTTGAAAGGATCGAAGAAAAGCTCGAAGAATCGTTCGAATTTCTTCGATTTTGACGCTGAGTTCGAAGCTGATTTCCAAGGTTTCAAAGATGATTCGTC
TATCGATTGCGATGATGATTTCGACGTCGGTGATGTTTTCGCCGATGTGAAACCATTCGTTTTCACTTCGACTCCAAAACCCGCCGTCTCCGCCGCTGCGGAAGGTAAATATTTAGGGATT
GACTTTAGTTTGAGCGATTGAATTGGTTAGAAATTAAGTTTCGATTCTATTTAGCTCTGTTCATAGATACCTTTATCTGCTTACGATTTTGATTGTTGTATAAAGCATGAGGTTTGATTTG
TGTGATATTTATATGCATATGAGGTTGAGTGATGAGACTTGATTTGTGTGATGAGATTTGATTAGAGATTGTTGATGAGTTATTTATAGAAGAACTCTTTTGTTTTGTTGTTGTTTCACTA
CTAGGTTCAGTTTTTGGTAAGAAAGTTACTGGCTTGGATGGGGACGCTGAGAAATCTGCAAATAGGAAGAGGAAGAATCAGTACCGAGGGATTAGGCAACGTCCTTGGGGAAAATGGGCTG
CTGAGATACGTGATCCAAGGGAAGGTGCTAGAATCTGGCTTGGAACGTTCAAGACAGCTGAGGAAGCTGCTAGAGCTTACGATGCTGCAGCGCGGAGAATCCGTGGATCTAAAGCTAAGGT
GAATTTCCCTGAAGAAAACATGAAGGCTAATTCTCAGAAACGCTCTGTGAAGGCTAATCTTCAGAAACCAGTGGCTAAACCTAACCCTAACCCAAGTCCAGCTTTGGTTCAGAACTCGAAC
ATCTCCTTTGAAAATATGTGTTTCATGGAGGAGAAACACCAAGTGAGCAACAACAACAACAACCAGTTTGGGATGACAAACTCCGTTGATGCTGGATGTAATGGGTATCAGTATTTCAGCT
CTGACCAGGGTAGTAATTCTTTCGATTGTTCGGAGTTTGGTTGGAGCGATCAAGCTCCGATAACTCCCGACATCTCTTCTGCGGTTATCAACAACAACAACTCAGCTCTGTTCTTTGAGGA
AGCCAATCCAGCTAAGAAGCTCAAGTCTATGGATTTCGAGACACCTTACAACAACACTGAATGGGACGCTTCACTGGATTTCCTCAACGAAGATGCTGTAACGACTCAGGACAATGGTGCA
AACCCTATGGACCTATGGAGTATTGATGAAATTCATTCCATGATTGGAGGAGTCTTC**TGA**AGAGATCCAGTTTCATGTAAATAAGGCTGCATGTTTGTGAGTTTCCCGCATCGTTCGTTTA
TCAACCTCCAAAACTTTCTAATGTCTGTTACTTGCATCTTCTTCTGCTGTCTCTGTCTGTCTCTCTCAGGAGTTCCTGTTTGCATTGCGAGAAGCCATGAGCCTCTATCTTGAGGGTAGTT
GTGATGAAGTTAAGTAGAGGCTTATTTTTAGGGGTTGTGGTAGTTTTTGTTTTAGTGAATCTTTTGAATTCGTTTGTGTTTTGTTTTTGTTACTTTATGCCCCAAAACTCCTTTAACATTT
GTCATAATGTGTTTGAACCTCTCATCTGTTTAATCAAATAAATCTTCTTTGTATGCTACTAAGAGTATGTGAGAACTGTTGAACATAACAAAGAAGTCACAAGCTAAGTTTGAACCAAACC
GGTTCTCGACGCGT

AtSLR1:D13RAP2.12

AAGCTTTAACATCCGGTACGTTAATTGCAATTGTAAATCCAGTCTCACATGAAAACCGGATAGAATTGGATCTAACCCAATGAACCGGGAATCTAAATTAAAAATTAAAATTGCTCTAATA
AGACATGTTAATCCTAGAATGCTAATCATGCTTCCGTTATGTTTTCCCTCCGTGAACACTAAATGGAGAATCATCAACAATATGTTAAGGATGTGTTTGTTTGGGCTGCTTCGGCAACTTT
CGCTGCTTGTTAGATTGCGACTTATTGCGAGCATAGATGTCATGGAAGTAGAATTCGAAATACATCGAGAAGATTAAAATACATCGAACGTAACGATAAATGCATTTACGAGTGCCATAGC
ATGACTTTCTTGGGGACGAGGATCATGGATGAATAAATATACTTGTTCAACAACATATGTTATGTTGCACGAGCAAATTAATTTCACATGGTGACGTTATCGACTAATGACAGTAGTTTGT
TGAAGTGAATCAATGAGGTACTGAAAAGTCATAGAACGTGGAAAATTACATTTGTCTTGTCTGCTAGAAATAATTGTAATGAGTTTTGCAAATTTGAAAACCAACATGTGAAGAAATCTAT
AAATATAGGTTTTAAGAAACAAAGAAGAGCATAAGAAAGAAAGTCGTGAAG**ATG**CCGAGGTCTCGCCGTGTTACTAGCGAGTTTATTTGGCCGGATCTGAAGAAGAATTTGAAAGGATCGA
AGAAAAGCTCGAAGAATCGTTCGAATTTCTTCGATTTTGACGCTGAGTTCGAAGCTGATTTCCAAGGTTTCAAAGATGATTCGTCTATCGATTGCGATGATGATTTCGACGTCGGTGATGT
TTTCGCCGATGTGAAACCATTCGTTTTCACTTCGACTCCAAAACCCGCCGTCTCCGCCGCTGCGGAAGGTAAATATTTAGGGATTGACTTTAGTTTGAGCGATTGAATTGGTTAGAAATTA
AGTTTCGATTCTATTTAGCTCTGTTCATAGATACCTTTATCTGCTTACGATTTTGATTGTTGTATAAAGCATGAGGTTTGATTTGTGTGATATTTATATGCATATGAGGTTGAGTGATGAG
ACTTGATTTGTGTGATGAGATTTGATTAGAGATTGTTGATGAGTTATTTATAGAAGAACTCTTTTGTTTTGTTGTTGTTTCACTACTAGGTTCAGTTTTTGGTAAGAAAGTTACTGGCTTG
GATGGGGACGCTGAGAAATCTGCAAATAGGAAGAGGAAGAATCAGTACCGAGGGATTAGGCAACGTCCTTGGGGAAAATGGGCTGCTGAGATACGTGATCCAAGGGAAGGTGCTAGAATCT
GGCTTGGAACGTTCAAGACAGCTGAGGAAGCTGCTAGAGCTTACGATGCTGCAGCGCGGAGAATCCGTGGATCTAAAGCTAAGGTGAATTTCCCTGAAGAAAACATGAAGGCTAATTCTCA
GAAACGCTCTGTGAAGGCTAATCTTCAGAAACCAGTGGCTAAACCTAACCCTAACCCAAGTCCAGCTTTGGTTCAGAACTCGAACATCTCCTTTGAAAATATGTGTTTCATGGAGGAGAAA
CACCAAGTGAGCAACAACAACAACAACCAGTTTGGGATGACAAACTCCGTTGATGCTGGATGTAATGGGTATCAGTATTTCAGCTCTGACCAGGGTAGTAATTCTTTCGATTGTTCGGAGT
TTGGTTGGAGCGATCAAGCTCCGATAACTCCCGACATCTCTTCTGCGGTTATCAACAACAACAACTCAGCTCTGTTCTTTGAGGAAGCCAATCCAGCTAAGAAGCTCAAGTCTATGGATTT
CGAGACACCTTACAACAACACTGAATGGGACGCTTCACTGGATTTCCTCAACGAAGATGCTGTAACGACTCAGGACAATGGTGCAAACCCTATGGACCTATGGAGTATTGATGAAATTCAT
TCCATGATTGGAGGAGTCTTC**TGA**AGAGATCCAGTTTCATGTAAATAAGGCTGCATGTTTGTGAGTTTCCCGCATCGTTCGTTTATCAACCTCCAAAACTTTCTAATGTCTGTTACTTGCA
TCTTCTTCTGCTGTCTCTGTCTGTCTCTCTCAGGAGTTCCTGTTTGCATTGCGAGAAGCCATGAGCCTCTATCTTGAGGGTAGTTGTGATGAAGTTAAGTAGAGGCTTATTTTTAGGGGTT
GTGGTAGTTTTTGTTTTAGTGAATCTTTTGAATTCGTTTGTGTTTTGTTTTTGTTACTTTATGCCCCAAAACTCCTTTAACATTTGTCATAATGTGTTTGAACCTCTCATCTGTTTAATCA
AATAAATCTTCTTTGTATGCTACTAAGAGTATGTGAGAACTGTTGAACATAACAAAGAAGTCACAAGCTAAGTTTGAACCAAACCGGTTCTCGACGCGT

**Supplementary Fig. S1**. Sequences for the SLR1 promoter-GUS and SLR1 promoter- RAP2.12 constructs. See methods for more information.

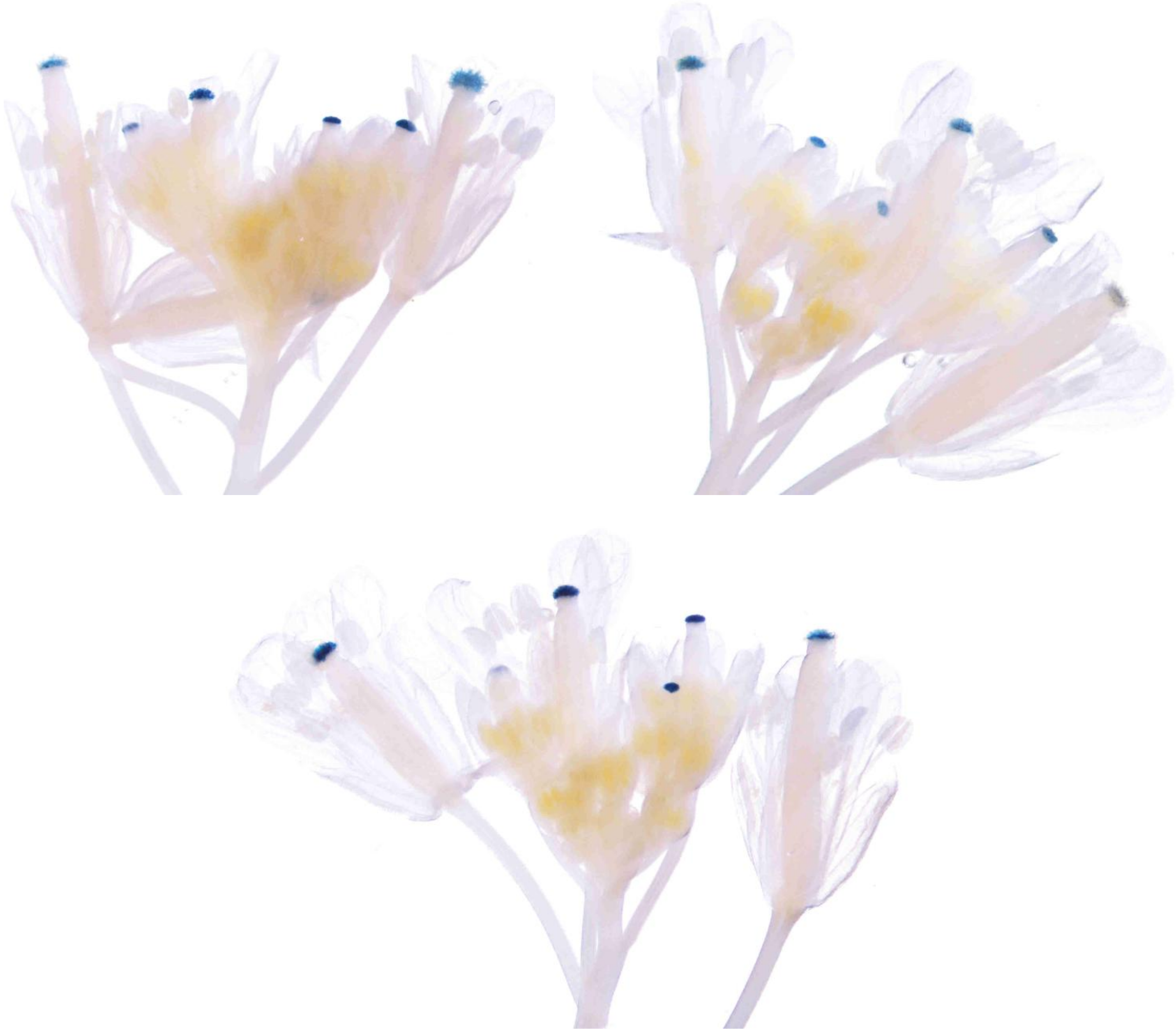

**Supplementary Fig. S2**. *A. thaliana* SLR1 promoter:GUS transgenic lines showing stigma-specific GUS activity in the inflorescences. Representative images from three different transgenic lines are shown.

For GUS staining, inflorescences were fixed in 80% acetone for overnight and then stained overnight at 37ºC in an X-Gluc solution (50 mM ferrocyanide, 50 mM ferricyanide, 500 mM $NaPO_4$, 20 mg/mL X-gluc). Stained pistils were mounted in water and imaged on a dissecting microscope.

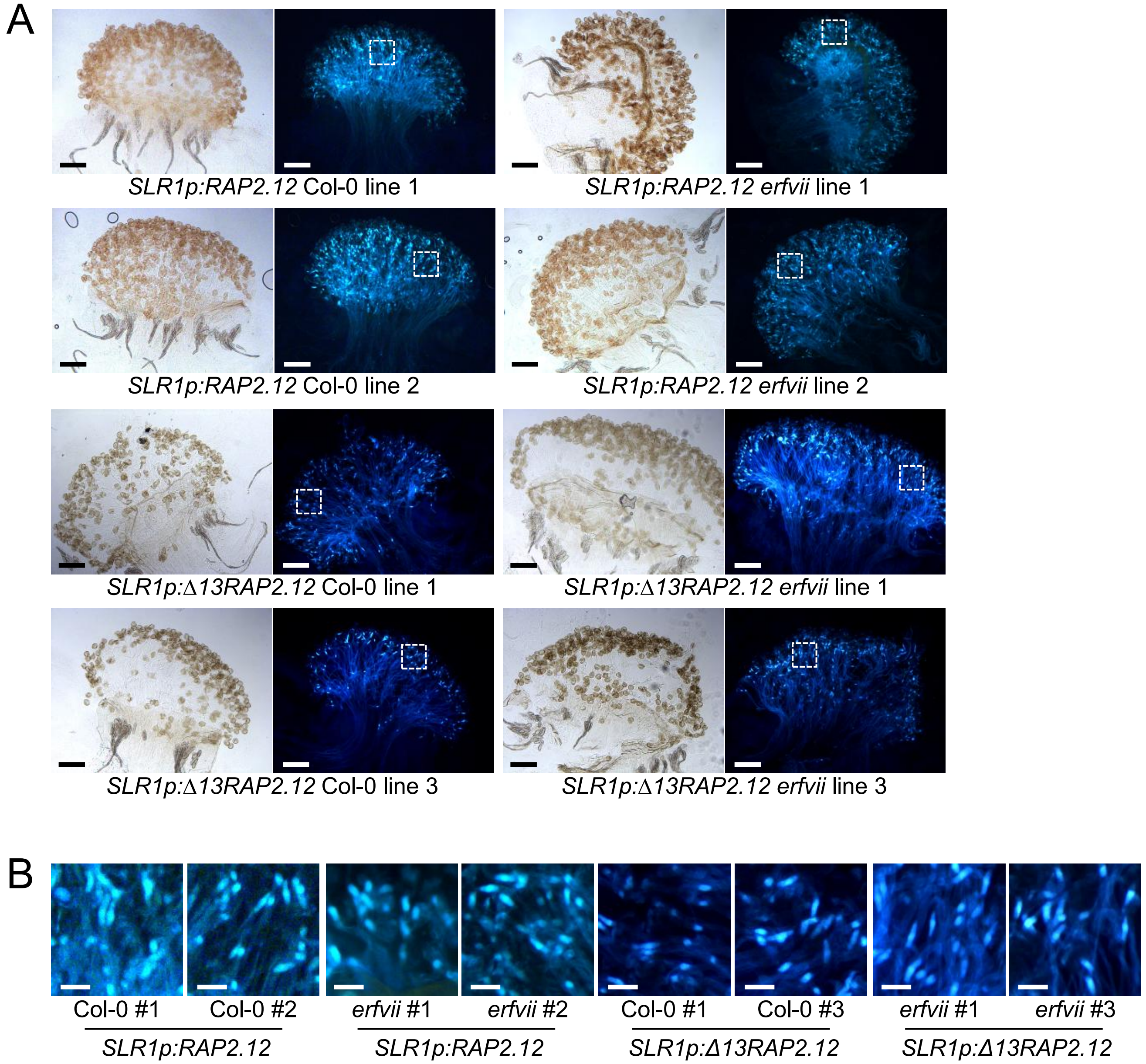


**Supplementary Fig. S3**. The stigma expression of *RAP2.12* and *Δ13RAP2.12* rescues the longer callose plug phenotype in wildtype Col-0 pollen tubes at 2 hours post-pollination.

(A) Additional representative images of wildtype Col-0 pollen tubes with callose plugs in stigmas from the *SLR1p:RAP2.12* and *SLR1p:Δ13RAP2.12* rescue lines at 2 hours post-pollination. Brightfield images (left) and aniline blue stained images (right) are shown for each sample. Scale bar = 100µm.

(B) Close up views of callose plugs in wildtype Col-0 pollen tubes growing in stigmas from the *SLR1p:RAP2.12* and *SLR1p:Δ13RAP2.12* rescue lines at 2 hours post-pollination. White boxes in (A) outline the regions shown here. Scale bar = 20µm.

*SLR1p:RAP2.12* Col-0 line 3 images are shown in Fig. 4, and lines 1 and 2 images are shown here.

*SLR1p:RAP2.12 erfvii* line 3 images are shown in Fig. 4, and lines 1 and 2 images are shown here.

*SLR1p:Δ13RAP2.12* Col-0 line 2 images are shown in Fig. 5, and lines 1 and 3 images are shown here.

*SLR1p:Δ13RAP2.12 erfvii* line 2 images are shown in Fig. 5, and lines 1 and 3 images are shown here.
